# The *App*^*NL-G-F*^ mouse retina is a site for preclinical Alzheimer’s disease diagnosis and research

**DOI:** 10.1101/2020.07.25.220707

**Authors:** Marjan Vandenabeele, Lien Veys, Sophie Lemmens, Xavier Hadoux, Géraldine Gelders, Luca Masin, Lutgarde Serneels, Jan Theunis, Takashi Saito, Takaomi C. Saido, Murali Jayapala, Patrick De Boever, Bart De Strooper, Ingeborg Stalmans, Peter van Wijngaarden, Lieve Moons, Lies De Groef

**Author notes:** Corresponding author: Lies De Groef, Neural Circuit Development and Regeneration Research Group, Department of Biology, University of Leuven (KU Leuven), Naamsestraat 61, box 2464, 3000 Leuven, Belgium.

## Abstract

In this study, we report the results of a comprehensive phenotyping of the retina of the *App*^*NL-G-F*^ mouse. We demonstrate that soluble Aβ accumulation is present in the retina of these mice early in life and progresses to Aβ plaque formation by midlife. This rising Aβ burden coincides with local microglia reactivity, astrogliosis, and abnormalities in retinal vein morphology. Electrophysiological recordings reveal signs of neuronal dysfunction yet no overt neurodegeneration was observed and visual performance outcomes were unaffected in the *App*^*NL-G-F*^ mouse. Furthermore, we show that hyperspectral imaging can be used to quantify retinal Aβ, underscoring its potential as a biomarker for AD diagnosis and monitoring. These findings suggest that the *App*^*NL-G-F*^ retina mimics the early, preclinical stages of AD, and, together with retinal imaging techniques, offers unique opportunities for drug discovery and fundamental research into preclinical AD.

## I. Introduction

Alzheimer’s disease (AD) is the number one neurodegenerative disorder and cause of dementia. Accumulation of amyloid-beta (Aβ) plaques and hyperphosphorylated Tau in tangles are two hallmarks of AD, and are believed to lead to neurotoxic inflammation, neuronal dysfunction, and eventually neurodegeneration. Despite decades of research, the AD field is still struggling with finding techniques for adequate diagnosis and disease monitoring, rational treatment strategies and valid research models.

AD is supposed to start 20 years before the first cognitive symptoms appear [20, 39, 51], and it is nowadays believed that one should focus on this time window for diagnosis and future treatment [19, 47]. In the search for new biomarkers and animal models that allow evaluation of this preclinical phase of AD, increasing attention has gone to the retina. Indeed, given that the retina is an integral part of the central nervous system, cellular and molecular mechanisms underlying neurodegeneration are conserved, and pathological processes occurring in the retina may be an indicator of those processes ongoing in other parts of the central nervous system [3, 11, 26]. In addition, state-of-the-art technologies for ocular imaging (e.g. optical coherence tomography and confocal scanning laser ophthalmoscopy) allow these processes to be visualised at a resolution of at least an order of a magnitude higher than conventional brain imaging techniques, without the need for invasive, costly procedures or tracers, and in a reproducible and quantifiable manner [11].

One emerging technique in the field of retinal biomarkers for AD, is retinal hyperspectral imaging. This non-invasive imaging spectroscopy technique relies on the wavelength-dependent effect of Aβ on light scatter, to measure the reflectance spectrum of a locus over a range of wavelengths and convert this into a hyperspectral score of which the magnitude correlates to the amount of Aβ. The exact pathological correlate of this hyperspectral signature is still unclear, although previous *in vitro* studies and theoretical modeling suggested that soluble Aβ_42_ is being measured [29, 30]. The latter would be a particular advantage, as converging evidence suggests that, in the preclinical phase of AD, soluble oligomeric Aβ accumulates to a threshold that eventually leads to symptomatic AD. Rather than intermediates to generation of Aβ plaques, these Aβ oligomers are nowadays regarded as being the most pathogenic form of Aβ. Aβ oligomers initiate neuropathological processes such as neuroinflammation, gliosis, degradation of synapses, impaired axonal transport, and oxidative damage, thereby leading to an enlarging pool of neurons with limited functionality and ultimately driving the diseased brain to the tipping point where actual neurodegeneration ensues [2, 7, 15, 52].

A multitude of studies has evaluated the retinal phenotype of AD mouse models, and revealed retinal abnormalities that mirror observations in human AD subjects, including elevated Aβ levels, Aβ deposits co-localizing with sites of apoptosis, tauopathy, neurodegeneration of ganglion and amacrine cells, microgliosis and astrogliosis, abnormal electrophysiology, as well as tight junction attenuation and vessel malformations [4, 10, 16, 24, 35]. However, results are often conflicting, and interpretation of their biological relevance is obscured by the fact that these transgenic AD mice all rely on ectopic overexpression of familial AD-associated genes (mutations). Here, we address this shortcoming with a comprehensive retinal phenotyping study of the *App*^*NL-G-F*^ knock-in mouse, which is characterized by physiological expression of humanized mouse APP, containing the Swedish, Beyreuther/Iberian and Arctic mutations, in disease-relevant central nervous system regions and cell types [32, 40]. For this first-time retinal phenotyping study of the *App*^*NL-G-F*^ mouse, we heavily relied on *in vivo* techniques to assess the possibility of longitudinal disease monitoring via retinal imaging and electrophysiology, besides the gold standard *post mortem* assays. Additionally, we evaluated the use of hyperspectral imaging to quantify retinal Aβ burden, based on the hypothesis that it detects soluble Aβ_42_ and therefore may offer a way to study preclinical AD. Altogether, this approach represents a novel tool box, with the retina of the *App*^NL-G-F^ mouse as a model and retinal imaging techniques as unique read-outs, for both fundamental and drug discovery research in the preclinical phase of AD.

## II. Materials and methods

### Animals

B6.129S5-*App*tm3.1Tcs (here referred to as *App*^*NL-G-F*^) [32, 40], *App*^*NL-G-F*^ x *CX*_*3*_*CR-1*^*GFP/+*^ (generated via crossing of *App*^*NL-G-F*^ and B6.129P2-*Cx3cr1*tm1Litt (The Jackson Laboratory)), B6;C3Tg (APPswe,PSEN1dE9)85Dbo/Mmjax (here referred to as APP/PS1) mouse lines and corresponding wild type (WT) controls were used. The *App*^*NL-G-F*^ mouse carries an *App* gene construct, which contains a humanized Aβ region and includes the Swedish “NL”, Iberian “F”, and Arctic “G” mutations. The APP/PS1 mouse contains transgenes for both APP and presenilin 1 (PSEN1 or PS1), under the control of the mouse prion promotor, with chimeric mouse/human APP with the Swedish mutation, and human PS1 lacking exon 9. The *App*^*NL-G-F*^ model was chosen as it avoids potential artifacts introduced by APP overexpression by using a knock-in approach to express APP at endogenous levels and with appropriate cell-type and temporal specificity. The APP/PS1 overexpression mouse was included in this study as a reference AD model, given that its retinal phenotype is well-described, e.g. [9, 12, 22, 29, 33, 45, 54]. Mice were bred under standard laboratory conditions. *App*^*NL-G-F*^ and WT female mice were sacrificed at 3, 6, 9, 12 and 18 months of age (± 1 month). Male mice were also included for the 18-month age group.

### Aβ extraction and ELISA detection

For soluble Aβ extraction, two retinas from one mouse were pooled and homogenized in lysis buffer (50 mM NaCl, 0.4 % diethanolamine, and protease inhibitor cocktail (Roche)). Samples were spun down and supernatant was centrifuged for 50 min at 100.000 x g. Supernatant containing the soluble Aβ fraction was collected. From the pellet, insoluble Aβ was extracted with GuHCl (6 M GuHCl/50 mM Tris-HCl) by sonication, incubation at 25°C for 60 min and centrifugation for 20 min at 170.000 x g. Aβ levels were determined via ELISA Meso Scale Discovery 96-well plates coated with antibodies for Aβ_40_ JRFcAβ40/28 and Aβ_42_ JRFcAβ42/26. JRFAβN/25 (human specific N-terminal epitope) labeled with sulfo-TAG was used as the detection antibody. Antibodies were provided by Dr. Marc Mercken (Janssen Pharmaceutica).

### Immunohistochemistry

Following perfusion with 4% paraformaldehyde (PFA), eyes were enucleated and post-fixated in 4% PFA for 1 hour. For cryosections (12 µm), eyes went through a sucrose gradient series for cryoprotection prior to embedding in O.C.T. compound (Tissue-Tek, Sakura). For wholemount preparations, retinas were flatmounted and post-fixated for one hour in 4% PFA. For Aβ immunostaining and hyperspectral imaging of retinal wholemounts, no additional fixation was done.

Retinal cryosections were stained for glial acidic fibrillary protein (GFAP), S100 calcium binding protein B (S100B), brain-specific homeobox/POU domain protein 3A (Brn3a), Ceh-10 Homeodomain-Containing Homolog (Chx10) or oligomeric Aβ (A11). For GFAP, antigen retrieval was performed via proteinase K treatment (5 min, 2 µg/ml, Qiagen). After blocking, sections were incubated overnight with anti-GFAP (1:2000, #Z0334, DAKO), anti-S100B (1:600, #ab52642, Abcam), anti-Brn3a (1:750, #sc-31984, Santa Cruz), anti-Chx10 (1:200, #X1180P, Sanbio) or A11 (1:500, #AHB0052, Invitrogen) primary antibodies. For GFAP, S100B, Brn3a and Chx10, an Alexa-conjugated secondary antibody (Invitrogen) was used; for A11, a biotin-conjugated antibody (Jackson ImmunoResearch) followed by tyramide signal amplification (Perkin Elmer) was used. 4′,6-diamidino-2-phenylindole (DAPI) was used as a nuclear counterstain.

For wholemount (immuno)staining, wholemounts were frozen for 15 min at −80°C, and incubated overnight with primary antibodies for melanopsin (1:5000, #AB-N38, Advanced Targeting Systems), ionized calcium-binding adapter molecule 1 (Iba1; 1:1000, #019-19741, Wako), choline acetyltransferase (ChAT; 1:500, #AB144P, Millipore), platelet-derived growth factor receptor beta (PDGFRβ) (1:500, #ab32570, Abcam), GFAP (1:1000, #Z0334, DAKO), RNA-binding protein with multiple splicing (RBPMS) (1:250, #1830-RBPMS, PhosphoSolutions), protein kinase C alpha (PKCα) (1:100, #sc-10800, Santa Cruz), green fluorescent protein (GFP; 1:500, #ab13970, Abcam) and/or Aβ (6E10, 1:500, #803001, BioLegend and 82E1, 1:300, #10323, Immuno-Biological Laboratories), or with biotinylated isolectin B4 (1:50, #B-1205, Vector Laboratories), followed by incubation with corresponding Alexa-conjugated secondary antibodies or Alexa-conjugated streptavidin (Invitrogen). For Aβ staining, a three-minute step in 100% formic acid was performed before freezing, as well as an extra blocking with goat anti-mouse Fab fragments (1:50, Jackson ImmunoResearch) for 1 hour.

### Microscopy and image analysis

Microscopy was performed either via confocal or multiphoton imaging (Olympus FV1000 or FV1000-M; for Aβ, Iba1, Cx3cr1-GFP, ChAT and isolectin B4) or conventional fluorescence microscopy (Leica DM6; for melanopsin, GFAP and S100B). For confocal or multiphoton imaging of retinal wholemounts, mosaic z-stack images (step size 2-4 μm) were taken. Image analyses were performed using Fiji software [42], unless stated otherwise.

i. *Microglia numbers and morphological parameters* were analyzed on maximum intensity projection images of retinal wholemounts, as described previously [5].
ii. *Macroglia reactivity* was quantified on retinal sections by manually counting GFAP-immunopositive fibers at the inner plexiform-inner nuclear layer border [43]. GFAP^+^ fibers were counted over a length of 300 µm, on 20 images per mouse (two peripheral and two central images per section; five sections per mouse, including the central section containing the optic nerve head, and the sections located 240 and 480 µm anterior/posterior). For S100B, the immunopositive area was measured in the nerve fiber/ganglion cell layer over a length of 300 µm, on 20 images per mouse (*cfr*. above) [18].
iii. *Retinal vasculature* was analyzed on retinal wholemounts, on maximum intensity projections from the upper vasculature. Vessel density and branching complexity was analyzed with AngioTool software [56] on a binarized image. Retinal vessel diameter was measured manually at a distance of 300-400 µm from the optic nerve head; 5 measurements were averaged per vessel.
iv. *RBPMS*^*+*^ *retinal ganglion cells* were counted on retinal wholemounts, using a validated automated counting method [27].
v. *Melanopsin*^*+*^ *retinal ganglion cell* numbers were counted manually on wholemount images [50].
vi. *ChAT*^*+*^ *cells* were counted in 3D, using the ‘spots’ module of Imaris software (version 7.4.2) on z-stack pictures of the wholemounts.

### Optical coherence tomography

Upon general anesthesia and pupil dilatation (0.5% tropicamide, Tropicol, Théa), optical coherence tomography scans of the retina (1000 A-scans, 100 B-scans, 1.4 x 1.4 mm, Bioptigen Envisu R2200) were acquired. Retinal layer thickness was measured using InVivoVue Diver 3.0.8 software (Bioptigen), at 16 locations equally spaced around the optic nerve head and averaged per mouse.

### Optomotor response test

Visual performance was tested using the virtual reality optomotor test (Optomotry, Cerebral Mechanics), as previously described [43]. Briefly, the mouse was place on a platform in the testing arena and, using the staircase method, the maximal spatial frequency of moving black-white stripes for which the optomotor reflex was still present, was determined.

### Electroretinogram

Following overnight dark adaptation, electroretinograms (Celeris, Diagnosys) were recorded for anesthetized mice. Lens electrodes with integrated stimulators were placed on the cornea after pupil dilation (0.5% tropicamide, Tropicol, Théa and 15% phenylephrine, Théa). Eyes were alternately stimulated, and the electrode on the contralateral eye was used as a reference. Full-field flash electroretinogram was recorded at increasing flash intensities of 0.003, 0.01, 0.1, 1, 2.5 and 7.5 cd*s/m^2^. For each intensity, 3 flashes were averaged. Inter-flash time increased from 5 s for the first two flash intensities, 10 s for the next two intensities to 30 and 45 s for the two last intensities. The scotopic threshold response was measured at 5*10^−6^ cd*s/m^2^ and 50 flashes were averaged. Analysis was performed with Espion v6.59.9 software (Diagnosys), as previously described [43]. The scotopic threshold response amplitude was defined as the peak of the curve. Using a band pass filter (75-300 Hz), the oscillatory potentials on the rising part of the b-wave were retrieved and the area under the curve was calculated. The latency of the oscillatory potentials was defined for OP2 [43].

### Hyperspectral imaging

A hyperspectral visible and near-infrared snapscan camera (150 bands, 470-900 nm, Imec; HSI Snapscan software (version 1.2.0)) was mounted on a Leica DM6 microscope with LED light source (400-700 nm). The snapscan camera acquires the hyperspectral image (2048 x 1536 pixels) by the movement of the linescan hyperspectral imaging sensor with on-chip filter technology on a miniaturized translation platform inside the camera body. Filters for each spectral band cover five adjacent pixel rows. The camera was used in stop-motion mode in steps of three pixel rows to reduce cross-talk between adjacent spectral bands and with HDR enabled (three frames) to reduce signal-to-noise ratio. Acquisition was set to cover only the 70 visible light spectral bands (471-691 nm). For hyperspectral imaging, retinal wholemounts were stained with DAPI, prior to mounting on a glass slide using a drop of saline and a cover slip. The transmission spectra of central (1 mm from the optic nerve head) and peripheral (2 mm from the optic nerve head) regions of each retinal quadrant were acquired using a 40x objective with the ganglion cell layer nuclei used as the focal plane.

Spectral analysis was performed using a modification of published methods [13] (Supplementary fig. 1). In brief, each frame of the recorded hyperspectral image was spatially resized to 20% of its original size to smooth out acquisition noise. Each spectrum was normalized by the average spectral value of the three longest wavelengths (685, 687 and 691 nm). Pixels with spectral values above 1.5 and below zero were considered as outliers and masked out from further analysis. Spectral data from all mice were stacked in a matrix and a corresponding class label (AD or wild type control [CO]) was given for each entry. The between class covariance spectrum was calculated. This spectrum corresponds to the average spectral difference between AD and CO groups. The hyperspectral score for a given spectral pixel was obtained by calculating the dot product between its spectral data and the between class covariance spectrum. The average spectral score for a given mouse was calculated using all pixels of all the HS images collected for that retina.

### Statistical analysis

Analyses were performed by a blinded observer. The employed statistical analyses and number of mice are stipulated in the respective figure legend. Analyses were performed using Prism v.8.2.1 (GraphPad). Differences were considered statistically significant for two-sided p-values < 0.05 (#, *p < 0.05; ##, **p < 0.01; ###, ***p < 0.001; ####, ****p < 0.0001; * for comparisons with baseline, # for comparisons between genotype).

## III. Results

### Amyloid accumulates in the retinas of *App*^*NL-G-F*^ mice, with Aβ plaque formation at old age

Retinal amyloid burden was studied in *App*^*NL-G-F*^ mice of various ages via ELISA. Soluble Aβ_42_ was detected at 3 months, with stable levels until 18 months, yet a steep increase at 24 months (p=0.0006) (Fig. 1a). Insoluble Aβ_42_, which is typically aggregated in amyloid plaques, was only clearly elevated at 12 months and dramatically increased at 24 months (p<0.0001) (Fig. 1a). In contrast, and in line with the combined effect of the Swedish mutation that increases total Aβ production and the Iberian mutation that increases the Aβ_42_/Aβ_40_ ratio, a steady level of soluble Aβ_40_ accumulation was present from 3 to 24 months, albeit two orders of magnitude lower than Aβ_42_ (Fig. 1b). Insoluble Aβ_40_ (Fig. 1b) as well as soluble/insoluble Aβ_38_ and Aβ_36_ levels (data not shown) were below the detection limit of the assay. For comparison, in the APP/PS1 overexpression mouse model, at 18 months of age (when retinal plaques are present [9, 12, 22, 33, 54]), we detected the accumulation of soluble Aβ_42_ (comparable levels to *App*^*NL-G-F*^), soluble Aβ_40_ (higher levels than *App*^*NL-G-F*^), and insoluble Aβ_42_ (lower levels than *App*^*NL-G-F*^) (Fig. 1c).

**Figure 1.**
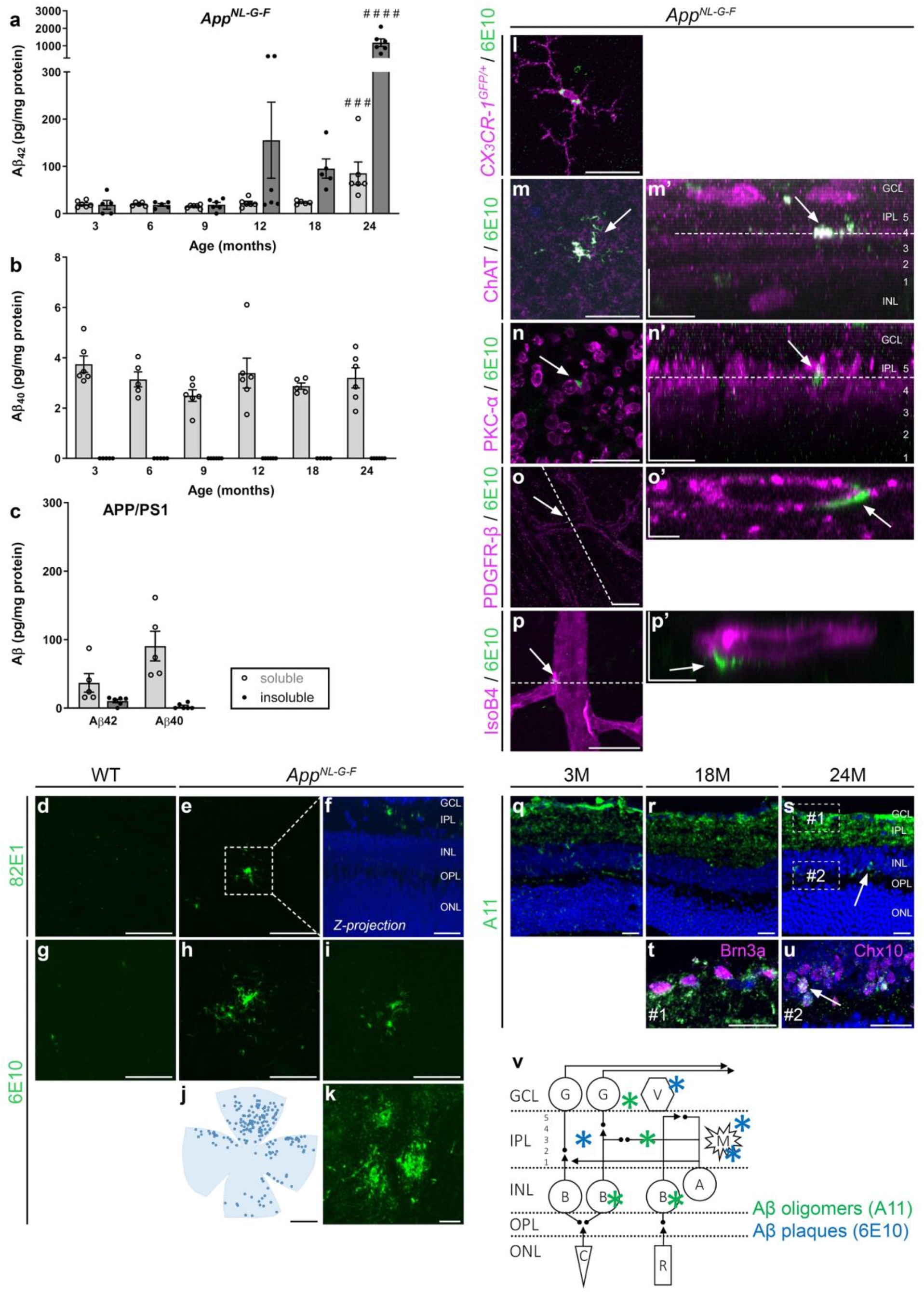
Amyloid burden increases with age in the retina of *App*^*NL-G-F*^ mice. (a-b) Amyloid ELISA on retinal lysates of *App*^*NL-G-F*^ mice shows accumulation of soluble Aβ_42_ from 3 months of age, which rises up till 24 months. Levels of insoluble Aβ_42_ only start to rise at 12 months. Accumulation of soluble, but not insoluble, Aβ_40_ is seen at all ages yet does not progress with age. One-way ANOVA with Dunnett’s multiple comparisons test (F_5,28_=6.686, p=0.0006 for soluble Aβ_42_; F_5,27_=20.27, p<0.0001 for insoluble Aβ_42_); n=5-6. (c) Eighteen-month-old APP/PS1 mice have comparable soluble Aβ_42_ levels yet lower levels of insoluble Aβ_42_ and higher soluble Aβ_40_ than *App*^*NL-G-F*^ mice of the same age. n=5. Of note, human Aβ levels are undetectable in the retinas of WT mice and therefore these controls are not shown. (d-i) Aβ immunostainings with antibodies 82E1 (d-f) and 6E10 (g-i) on retinal wholemounts of 18-month-old WT (d, g) and *App*^*NL-G-F*^ mice (e, f, h, i) reveal the presence of Aβ plaques in *App*^*NL-G-F*^ mice only. (f) Example of a reconstruction of the z-plane of the image in panel e, with 82E1 and DAPI nuclear counterstaining, showing that Aβ plaques mainly localize to the inner plexiform layer. (j) Representative example of a retinal wholemount on which 6E10-positive Aβ plaques are depicted with dots to illustrate plaque distribution in the retina. (k) Aβ plaques in the cerebral cortex of an 18-month-old *App*^*NL-G-F*^ mouse. (l-p) Double labeling for Aβ (6E10) and retinal cell markers on retinal wholemounts of 18-month-old *App*^*NL-G-F*^ mice. Single-tile images of the retinal flatmounts, with corresponding orthogonal projections of the z-stack (‘) are shown. Dotted lines in m’ and n’ indicate the z depth for with the single-tile image is shown. Dotted lines in o and p indicate the orientation of the orthogonal projections shown in o’ and p’. (l) Double labeling for Aβ and Iba1 shows both extra- and intracellular Aβ deposits in microglia, suggesting that microglia surround and phagocytose plaques. (m-n) Most plaques are found in the inner plexiform layer, where they align with the ChAT^+^ strata in which the terminals of the cholinergic amacrine cells are organized (m) and the PKCα^+^ sublayer in which the terminals of the bipolar cells are organized (n). (o-p) Plaques are also found in the uppermost layers of the retina, in association with the pericytes (m) and endothelial cells (n) of the retinal blood vessels. (q-s) Immunostaining for oligomeric amyloid (A11) on retinal cross-sections of 3-month- (3M) (q), 18-month- (18M) (r) and 24-month-old (24M) *App*^*NL-G-F*^ mice (s) shows progressive accumulation of Aβ oligomers. (t, u) These Aβ oligomers are mostly found in the inner plexiform layer, in association with ganglion cells (Brn3a^+^) and displaced amacrine cells (Brn3a^−^) in the ganglion cell layer (t), and inside Chx10^+^ bipolar cells (u). (v) Schematic overview of the retina and the locations in which Aβ oligomers and plaques were found. Scalebars: 10 µm (m’-p’), 20 µm (d-i, k, n, q-u), 50 µm (p), 100 µm (l, m, o), 1 mm (j). Data are shown as mean ± SEM. GCL = ganglion cell layer, IPL = inner plexiform layer, INL = inner nuclear layer, OPL = outer plexiform layer, ONL = outer nuclear layer, G = ganglion cell, A = amacrine cell, B = bipolar cell, V = blood vessel, M = microglia, C = cone, R = rod.

In accordance with these results, Aβ plaques were detected via 6E10 and 82E1 immunohistochemistry on retinal wholemounts of *App*^*NL-G-F*^ mice of 18 months of age (Fig. 1d-i). Morphologically, these resembled classical brain plaques, with a dense core and radiating fibrillary arms (Fig. 1k). Sizes typically ranged from 5 to 20 µm. Aβ plaque density was higher in the peripheral than in the central retina, while no overt differences between the retinal quadrants were seen (Fig. 1j). Plaques mainly localized in the inner plexiform layer (Fig. 1f), often closely associated with microglia (Fig. 1l & Fig. 4d-g), ChAT^+^ strata with cholinergic amacrine cell synapses (Fig. 1m), and PKCα^+^ strata with bipolar cell terminals (Fig. 1n). In addition, some were found around the blood vessels in the nerve fiber layer, in close association with pericytes (Fig. 1o) and endothelial cells (Fig. 1p). In all these cases, the Aβ deposits were extracellularly. Intracellular 6E10 staining was only seen in microglia (Fig. 1l), probably due to phagocytosis of Aβ plaques; and in RBPMS^+^ and melanopsin^+^ ganglion cells (data not shown), likely reflecting endogenous APP expression. Plaques did not associate with ganglion cells (RBPMS and melanopsin), bipolar (Chx10) and amacrine (ChAT) cell somata, nor astrocytes and radial Müller glia (GFAP) (data not shown). Oligomeric Aβ was visualized with an A11 immunostaining on retinal sections and was most abundant in all strata of the inner plexiform layer – comprising amacrine and bipolar cell presynaptic and ganglion cell postsynaptic terminals (Fig. 1q-u). Oligomeric Aβ was also observed extracellularly in the ganglion cell layer (Fig. 1t) and intracellularly in Chx10^+^ bipolar cells (Fig. 1u). A progressive accumulation over time – in 3- to 18- to 24-month-old mice –, increasingly affecting the inner plexiform layer and bipolar cells, was observed (Fig. 1q-s). No Aβ staining, with 6E10 nor A11, was observed in the outer retina.

These data show that, while soluble Aβ_42_ (and Aβ_40_) is present in the retinas of *App*^*NL-G-F*^ mice early in life, retinal Aβ plaques only appear from 18 months of age onwards. Given this relatively slow progression of amyloidopathy, this retinal mouse model may be used to study the conversion from early to late stages of AD pathogenesis, as well as the effect of soluble Aβ on retinal structure and function. For the characterization of the latter, in addition to longitudinal studies, we will largely focus on ages 3 and 18 months in the rest of this study, as these represent disease stages without and with Aβ plaques in the retina, respectively.

### Hyperspectral imaging detects early retinal accumulation of Aβ in *App*^*NL-G-F*^ mice

Aiming to provide a proof-of-concept for the use of hyperspectral imaging for the early detection of retinal Aβ accumulation and for monitoring its progression, we next performed *ex vivo* hyperspectral imaging of retinal wholemounts of 3- and 18-month-old *App*^*NL-G-F*^ mice and age-matched WT controls. For both ages, the average spectra of the *App*^*NL-G-F*^ mice were clearly distinct from WT. In young mice, this separation was most marked in the 475-500 nm range (Fig. 2a), while in old mice the spectral separation became even more apparent, extending to 600 nm (Fig. 2b). A hyperspectral score was calculated for each mouse from the *ex vivo* retinal reflectance data, using a modification of methods developed for *in vivo* hyperspectral retinal imaging [13]. Hyperspectral scores were significantly higher in *App*^*NL-G-F*^ mice compared to age-matched WT mice at both ages (p=0.0014 at 3 months, p=0.0002 at 18 months) (Fig. 2d-e). Furthermore, the difference in the mean hyperspectral score of *App*^*NL-G-F*^ *versus* age-matched WT mice was also higher at 18 months than at 3 months, in line with the progressive accumulation of Aβ seen in the ELISA. Hyperspectral scores were calculated for each retinal quadrant in both central and peripheral locations (Fig. 2j). In keeping with the immunohistochemistry findings, which demonstrated a predominately peripheral distribution of Aβ, hyperspectral scores were higher in the peripheral retina than in the central retina (Fig. 2g-h).

**Figure 2.**
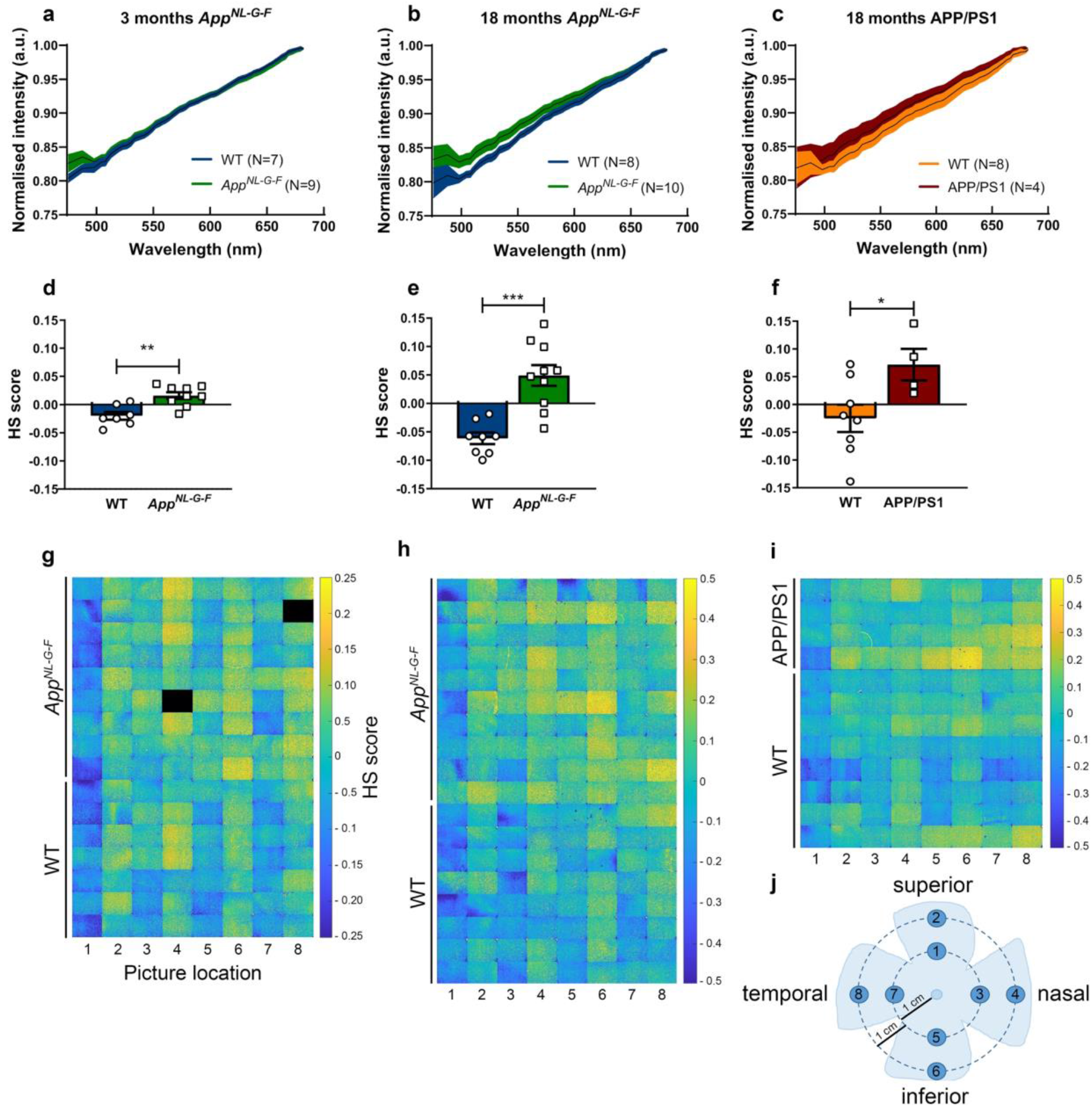
Quantification of retinal Aβ burden via hyperspectral imaging reveals progressive retinal amyloidosis in *App*^*NL-G-F*^ mice. (a) Average spectra of 3-month-old *App*^*NL-G-F*^ and WT mice are different, with a clear separation in the short wavelength range. (b) Average spectra of 18-month-old *App*^*NL-G-F*^ and WT mice show significant spectral differences for all wavelengths up to 600 nm. (c) Average spectra of 18-month-old APP/PS1 and WT mice show significant spectral differences from ∼500 to ∼600 nm. Data are shown as mean ± 95% confidence intervals. (d-e) For both ages, comparison of hyperspectral (HS) scores (obtained by compression of the spectral information to maximize between group separation) of *App*^*NL-G- F*^ and WT mice shows a strong effect size and significant difference between genotypes. Unpaired two-tailed t-tests (t_14_=3.957,p=0.0014 at 3 months; t_16_=4.919,p=0.0002 at 18 months); n=8-10. (f) HS scores of 18-month-old APP/PS1 and WT mice are significantly different. Unpaired two-tailed t-tests (t_10_=2.395,p=0.0376); n=4-8. (g-i) Heatmaps with a spatial representation of the HS score per image for each mouse. Note that image fields 2, 4, 6 and 8, which correspond to the peripheral retinal locations, have higher scores, indicative for a higher amyloid load. (j) Schematic representation of the retinal imaging locations.

Remarkably, 18-month-old APP/PS1 mice had retinal hyperspectral signatures that closely mimics those of 18-month-old *App*^*NL-G-F*^ mice (Fig. 2c, f, i). Given that these mice have similar levels of soluble Aβ_42_, but markedly different levels of soluble Aβ_40_ and insoluble Aβ_42_, it is possible that soluble Aβ_42_ is most contributory to the hyperspectral signature, as previously suggested by Vince and More [29].

### Neuronal dysfunction yet no degeneration in the inner retina of *App*^*NL-G-F*^ mice

Many studies have shown neurotoxic effects of soluble Aβ, and that functional deficits may precede neurodegeneration. Therefore, first electroretinography was performed with *App*^*NL-G-F*^ and WT mice at different time points. Overall, no abnormalities were observed at early ages (Supplementary fig. 2). At 18 months, however, the latency of both the oscillatory potentials and the b-wave, a measure for amacrine and bipolar cell function, respectively, showed significant changes (p<0.0001 for the effect of genotype) in the *App*^*NL-G-F*^ mice relative to age-matched WT mice (Fig. 3d-g). Furthermore, also at 18 months of age, the latency of the a-wave was affected at the lowest light intensity (p<0.0001), suggesting compromised photoreceptor function (Fig. 3b-c).

**Figure 3.**
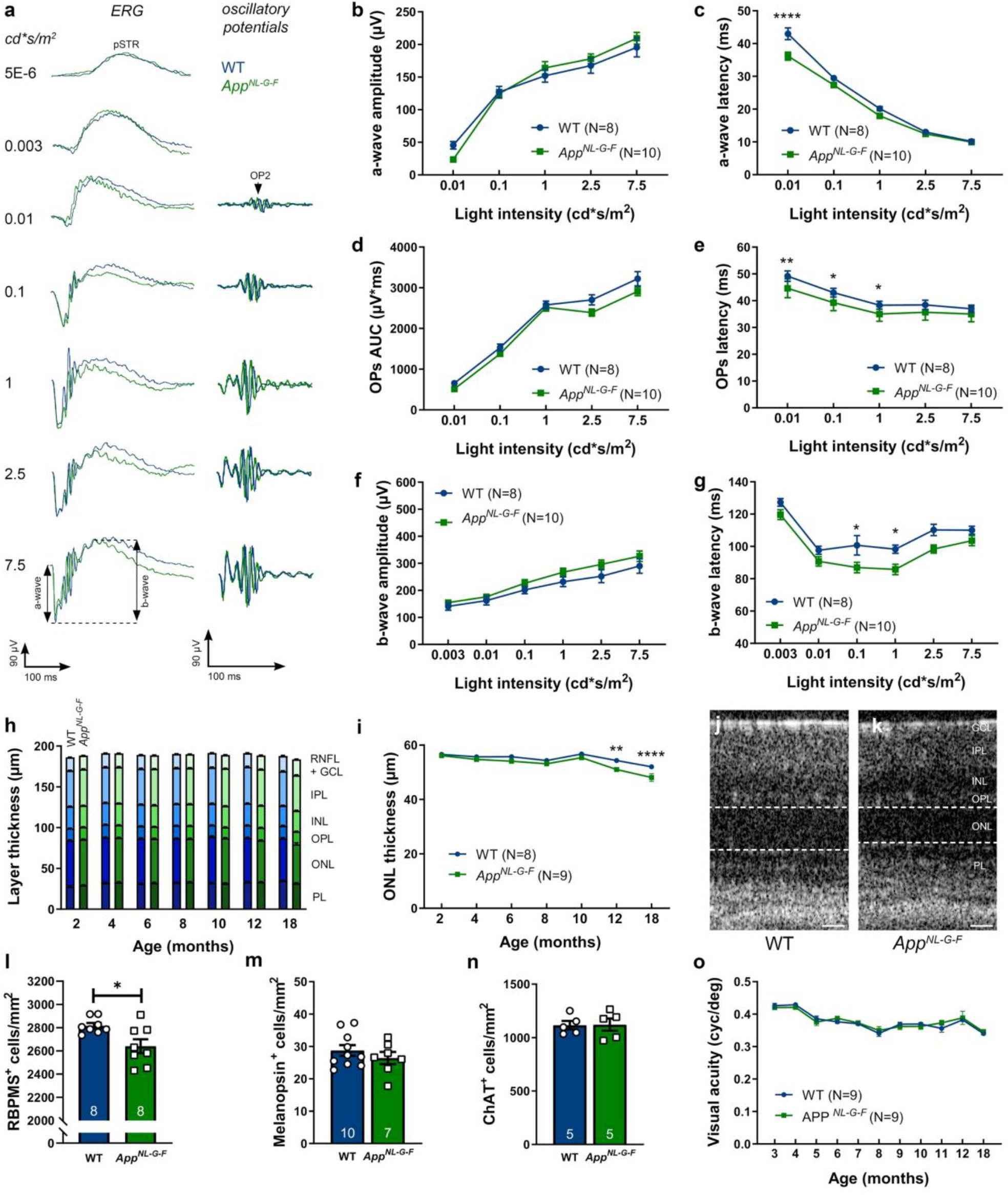
Neuronal dysfunction yet no degeneration in the retina of *App*^*NL-G-F*^ mice. (a) Representative electroretinogram measurements, with increasing light intensities, in 18-month-old *App*^*NL-G-F*^ (green) and WT (blue) mice. (b) The amplitude of the a-wave is unaffected in *App*^*NL-G-F*^ mice of 18 months, yet (c) the latency time is decreased for the lowest light intensity. Two-way ANOVA with Sidak’s multiple comparisons test (F_1,85_ = 23.14 for the effect of genotype, p<0.0001); n=8-10. (d) Oscillatory potentials show a similar wave front (measured as the area under the curve, AUC), but (e) a reduced latency in 18-month-old *App*^*NL-G-F*^ mice, compared to WT. Two-way ANOVA with Sidak’s multiple comparisons test (F_1,80_= 37.14 for the effect of genotype, p=0.0013, 0.0124, 0.0378 for 0.01, 0.1 and 1 cd*s/m^2^ respectively); n=8-10. (f) Similarly, the amplitude of the b-wave is unaltered, but (g) the latency time is decreased in 18-month-old *App*^*NL-G-F*^ mice compared to WT mice. Two-way ANOVA with Sidak’s multiple comparisons test (F_1,97_ = 27.90 for the effect of genotype, p=0.0206 and 0.044 for 0.1 and 1 cd*s/m^2^); n=8-10. Full electroretinogram data is given in Supplementary figure 2. (h) Analysis of the thickness of the retinal layers, as measured by optical coherence tomography, shows no differences between *App*^*NL-G-F*^ and WT mice from 2 to 18 months of age, (i) with the exception of a thinning of the outer nuclear layer. Two-way ANOVA with Sidak’s multiple comparisons test (F_1,106_ = 32.14 for the effect of genotype, p=0.0016 and p<0.0001 for 12 and 18 months, respectively); n=8-9. (j-k) Representative optical coherence tomography images of WT (j) and *App*^*NL-G-F*^ (k) retinas. Dotted lines delineate the outer nuclear layer. (l) Quantification of RBPMS^+^ retinal ganglion cell density reveals ganglion cell loss in 18-month-old *App*^*NL-G-F*^ compared to WT mice. Unpaired two-tailed t-test (t_14_=2.745,p=0.0158); n=8. (m-n) Cell counts of melanopsin^+^ ganglion cells (m) and ChAT^+^ neurons (n) show no differences in numbers between 18-month-old *App*^*NL-G-F*^ and WT mice. (o) A longitudinal study of visual acuity in *App*^*NL-G-F*^ mice from 3 till 18 months revealed a decrease with age but no genotype differences. Two-way ANOVA (F_10,180_=21.11 for the effect of age, F_1,180_=0.1149 for the effect of genotype); n=9. Scale bar: 25 µm. Data are depicted as mean ± SEM; RNFL = retinal nerve fiber layer, GCL = ganglion cell layer, IPL = inner plexiform layer, INL = inner nuclear layer, OPL = outer plexiform layer, ONL = outer nuclear layer, PL = photoreceptor layer.

Next, we assessed potential neurodegeneration in the *App*^*NL-G-F*^ retina via a longitudinal *in vivo* optical coherence tomography experiment. No differences in the overall thickness of the retina nor the nerve fiber, ganglion cell, inner plexiform or inner nuclear layers were identified at any age (Fig. 3h). Of note, no neurodegeneration was observed in the brains of the *App*^*NL-G-F*^ mice either [40]. A small decrease in the thickness of the outer nuclear layer was found in *App*^*NL-G-F*^ mice, from 12 months (p=0.0016 for 12 months, p<0.0001 for 18 months) of age onwards (Fig. 3i). Although these findings corroborate the electroretinogram results, we are unsure how to interpret their biological relevance given that this thinning is limited and occurs in the outer retina – where plaques nor oligomeric Aβ were seen. Loss or dysfunction of the retinal pigment epithelium due to amyloidopathy in these support cells may underlie this photoreceptor degeneration [6, 34], or, alternatively, the AD phenotype of the *App*^*NL-G-F*^ mice may lead to accelerated aging and therefore earlier age-related deterioration/loss of rods [21]. This should be further investigated in follow-up studies.

Finally, based on the localization of plaques in the nerve fiber and inner plexiform layers, and in order to confirm and complement the optical coherence tomography data, we next examined neuronal subpopulations in the inner retinas of 18-month-old animals. We quantified: (i) ganglion cells, using the pan-ganglion cell marker RBPMS (Fig. 3l); (ii) melanopsin^+^ ganglion cells, given that this subtype appears to be selectively vulnerable to Aβ pathology [31] (Fig. 3m); and (iii) ChAT^+^ cholingeric amacrine cells (Fig. 3n). While a small loss of RBPMS^+^ ganglion cells was observed (6.26 ± 2.28%, p=0.0158), melanopsin^+^ and ChAT^+^ neuronal populations did not show any sign of degeneration in 18-month-old *App*^*NL-G-F*^ mice. In line with all of the above observations, and in keeping with the relatively minor loss of ganglion cells and neural plasticity leading to reversible/inducible changes of retinal ganglion cell electrical activity during a critical period of dysfunction preceding death [37], 18-month-old *App*^*NL-G-F*^ mice behaved normally in the optomotor response test for visual acuity (Fig. 3o) and contrast sensitivity (data not shown). Altogether, these results indicate that the majority of inner retinal neurons in the *App*^*NL-G-F*^ mouse survive progressive amyloidosis up till 18 months. Nevertheless, although no cell death was observed at this age, amacrine and bipolar cell function may be used as a sensitive read-out for AD pathology in the retina of *App*^*NL-G-F*^ mice.

### Local microglia reactivity, astrogliosis and vascular changes in the retina of aged *App*^*NL-G-F*^ mice

Because of the strong implication of microgliosis and astrogliosis in AD pathogenesis, and as neurotoxic inflammation may precede neurodegeneration, we next examined the retinal glial cells. First, microgliosis was investigated in entire Iba1-labeled retinal wholemounts of *App*^*NL-G-F*^ mice aged between 3 and 18 months. Overall quantification of microglial density (Fig. 4a), soma size (Fig. 4b) and roundness (data not shown) did not reveal population-wide differences between the two genotypes at any age. Analysis of 18-month-old *App*^*NL-G-F*^ x *CX*_*3*_*CR-1*^*GFP/+*^ mice retinas, with GFP-labeled microglia, confirmed these findings (data not shown). However, a detailed investigation of these retinas, co-labeled for 6E10, revealed local microglia reactivity around Aβ plaques, as was previously shown in the brains of these mice [40] and in the retinas of other AD mouse models (3xTgAD, APP/PS1) [10, 36, 41]. Indeed, microglia clustered around Aβ plaques, and these cells showed less ramified, thicker processes and larger somas (p=0.0112) compared to microglia in regions without plaques (Fig. 4c-g) or in wild type *CX*_*3*_*CR-1*^*GFP/+*^ mice. Second, Müller glia and astrocyte reactivity were assessed on transverse sections of the retina. Longitudinal assessment of Müller glia reactivity, by counting the number of GFAP-positive radial fibers, revealed higher reactivity in 18-month-old *App*^*NL-G-F*^ *versus* WT mice (p=0.0432) (Fig. 4h-j). Further substantiating these findings, and in line with the brain phenotype [40], analysis of the S100B immunopositive area, which is specific for astrocytes, showed astrogliosis in the retina of 18-month-old *App*^*NL-G-F*^ mice (p=0.0057) (Fig. 4 k-m).

**Figure 4.**
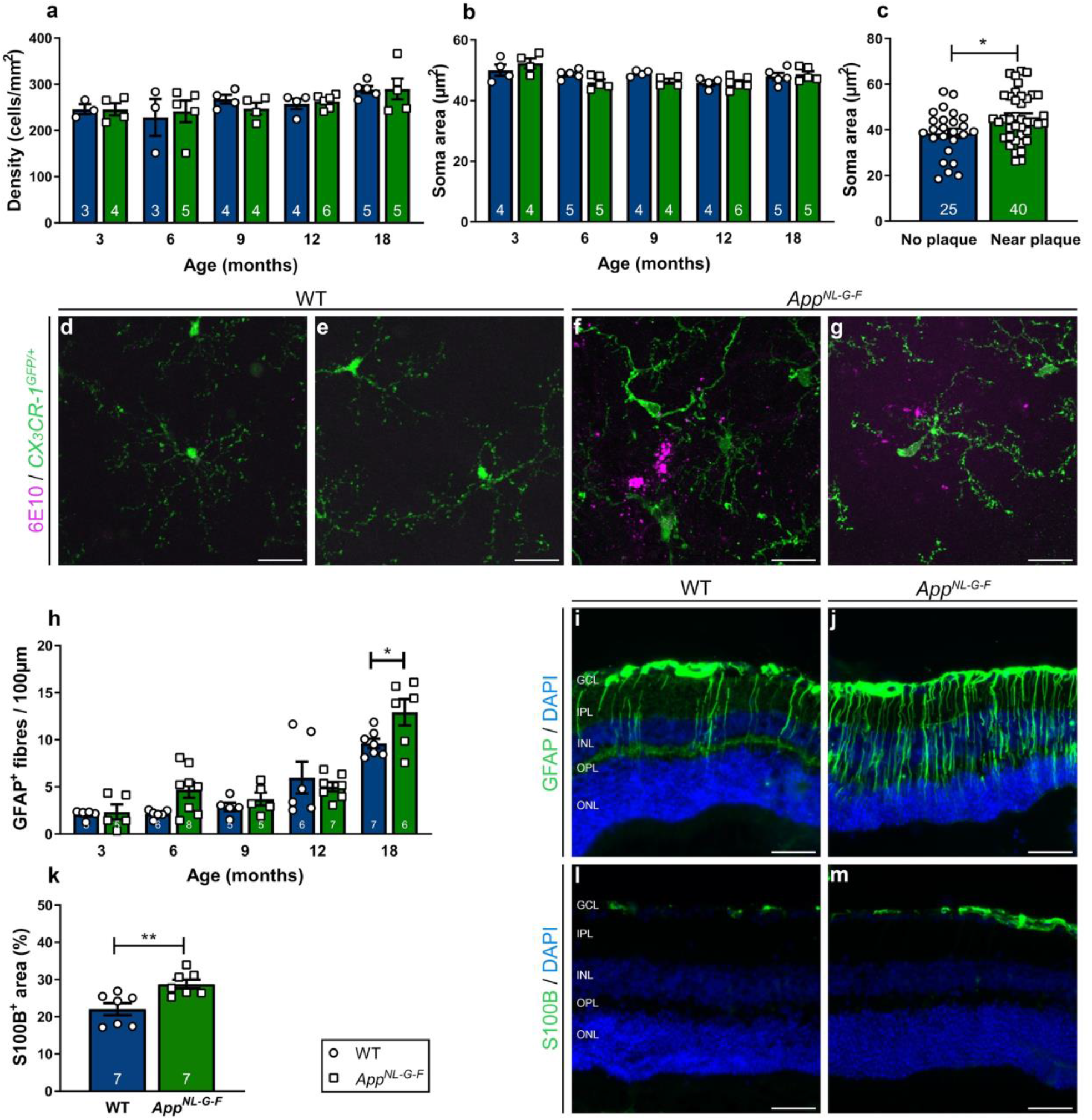
Locally reactivated microglia and macrogliosis in the retinas of aged *App*^*NL-G-F*^ mice. (a-b) Quantification of microglial density (a) and soma area (b) on Iba1-labeled retinal wholemounts of 3-to 18-month-old *App*^*NL-G-F*^ and WT mice reveals no differences between genotypes when analyzed for the entire retina. An effect of aging was observed for microglia density (Two-way ANOVA, F_4,33_ = 2.976, p=0.0334); n=3-6. (c) Morphometric analysis of plaque-associated microglia *versus* microglia distant from Aβ plaques in 18-month-old *App*^*NL-G-F*^ x *CX*_*3*_*CR-1*^*GFP/+*^ retinas reveals that plaque-associated microglia have a larger soma, which is indicative of their activation. Unpaired two-tailed t-test (t_63_=2.614, p=0.0112); n=25-40 cells from 6 mice. (d-g) Representative images of immunostaining for Aβ (6E10) on retinal wholemounts of 18-month-old *App*^*NL-G-F*^ x *CX*_*3*_*CR-1*^*GFP/+*^ (f-g) and *CX*_*3*_*CR-1*^*GFP/+*^ mice (d-e), illustrating that green fluorescent microglia surrounding Aβ plaques display morphological alterations typical for reactive microglia, with thicker and less ramified processes and a larger soma. (h) Counting the number of GFAP^+^ radial fibers on retinal sections of *App*^*NL-G-F*^ and WT mice shows that macrogliosis increases with age, and significantly differs between genotypes at 18 months. Two-way ANOVA with Sidak’s multiple comparisons test (F_4,50_=34.02 for the effect of age, F_1,50_=4.411 for the effect of genotype, p=0.0432); n=5-8. (i-j) Representative images of a GFAP immunostaining on retinal cross-sections of 18-month-old WT (i) and *App*^*NL-G-F*^ (j) mice. (k) Quantification of the S100B immunopositive area on retinal sections of 18-month-old *App*^*NL-G-F*^ and WT mice shows astrogliosis in the *App*^*NL-G-F*^ retina. Unpaired two-tailed t-test (t_12_=3.358, p=0.0057); n=7. (l-m) Representative images of the S100B immunostaining on WT (l) and *App*^*NL-G-F*^ (m) retinas. Scalebar (d-g): 20 µm, scalebar (i, j, l, m): 50 µm. Data are shown as mean ± SEM. GCL = ganglion cell layer, IPL = inner plexiform layer, INL = inner nuclear layer, OPL = outer plexiform layer, ONL = outer nuclear layer.

Looking at non-neuronal cell types, besides glial cells, also vasculature may undergo pathological alterations in the context of AD. Changes in retinal blood vessel function and morphology, including branching complexity, vessel density and vessel diameter, are being explored as potential early biomarkers for AD [8, 38, 53, 55]. Given that a subset of plaques was found closely associated with the retinal vasculature, we next assessed vessel morphology on retinal wholemounts stained with isolectin B4 (Fig. 5). Vessel density (Fig. 5a) and branching complexity (Fig. 5b) were unaltered in *App*^*NL-G-F*^ *versus* WT mice at 18 months of age. However, diameters of the retinal venules (Fig. 5c), yet not of the retinal arterioles (Fig. 5d), were smaller in *App*^*NL-G-F*^ mice (p=0.0235). Of note, Shi *et al*. recently reported pericyte loss accompanying retinal vascular amyloidosis, hence the observed reduction in diameter may be a manifestation of this [44].

**Figure 5.**
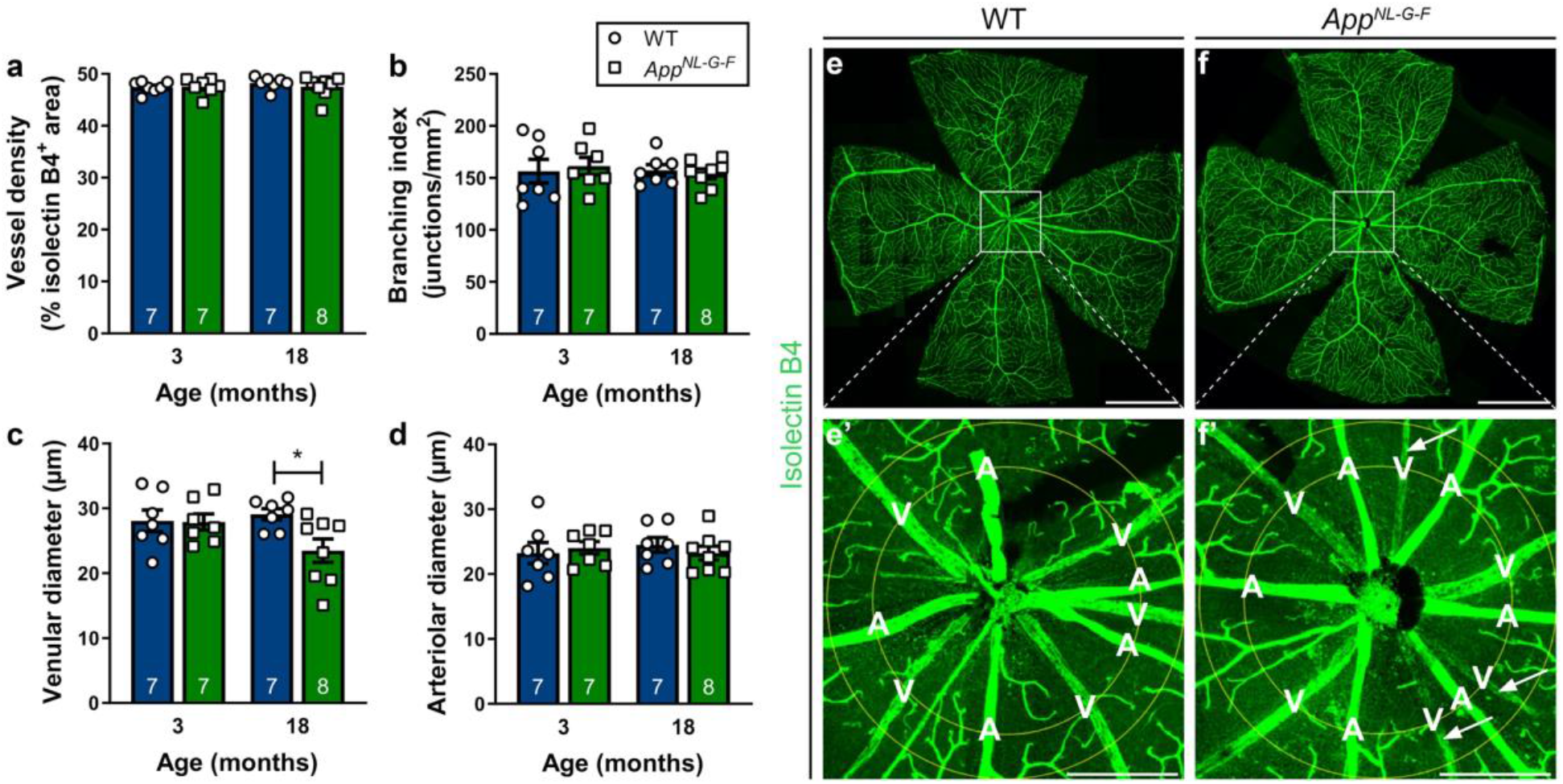
Retinal venules of *App*^*NL-G-F*^ mice have reduced diameters. (a-b) Retinal vasculature was analyzed at 3 and 18 months of age. Quantification of the density of the vascular network (a) and its branching complexity (b) shows no differences according to genotype. (c-d) (c) Retinal venular diameters are reduced in 18-month-old *App*^*NL-G-F*^ mice. Two-way ANOVA with Sidak’s multiple comparison’s test (F_1,25_=3.807 for the effect of genotype, p=0.0235); n=7 for 3 to 6 months, n=7-8 for 18 months. (d) No differences are seen in retinal arteriolar diameters. (e-f) Representative images of isolectin B4-labeled retinal flatmounts of 18-month-old WT (e) and *App*^*NL-G-F*^ (f) mice. (e’-f’) Magnification of the region of interest in which vessel analysis was performed. Arrows indicate retinal venules with a reduced diameter. Scalebar: 1000 µm. Data are depicted as mean ± SEM. A = artery, V = vein.

## IV. Discussion

In this study, we investigated the extent to which AD-associated disease processes are present in the retina of the *App*^*NL-G-F*^ knock-in mouse, and whether they can be monitored via *in vivo* retinal imaging and electrophysiology. First, we demonstrated that soluble Aβ (mostly Aβ_42_) accumulation is present early in life in these mice yet retinal Aβ plaques are only observable at 12 months of age and older. This rising Aβ burden in the retina coincides with local microglia reactivity, astrogliosis, and abnormalities in retinal vein morphology, and leads to compromised electrophysiological function of selected retinal neurons. No overt neurodegeneration was seen, however, and visual performance outcomes were intact. Altogether, these findings suggest that the *App*^*NL-G-F*^ retina resemble changes that occur in the brain in preclinical AD, during which Aβ accumulation instigates the earliest detrimental effects on astrocytes, microglia and vasculature, poses a stress on neuronal functionality but does not lead to overt functional impairment and neuronal death yet. Second, the data presented in this study suggest that there is an identifiable pre-plaque stage in AD in which only soluble forms of Aβ are present, followed by a phase in which soluble, oligomeric forms of Aβ are likely in equilibrium with plaques. The *App*^*NL-G-F*^ retina and reported retinal imaging techniques offer unique opportunities for further fundamental and preclinical research into this sequence of biological events over the course of preclinical AD. Hence, the *App*^*NL-G-F*^ mouse is not only the first AD model to accumulate Aβ without phenotypes related to APP overexpression and to more closely model human pathophysiology, its retina also is the first model organ to study preclinical AD. Combined with the available non-invasive, high-resolution imaging techniques and functional assessments of the retina, this creates a toolbox for researchers to shed new light on Aβ proteostasis and novel disease-modifying therapies focused thereon.

The rather mild phenotype is in stark contrast to observations made in the brains of *App*^*NL-G-F*^ mice. Indeed, previous studies have shown that (sub)cortical Aβ plaques appear at 2-4 months of age, with astrogliosis and microglial reactivity at 6-9 months, and cognitive impairment as early as 6 months [40]. The later formation of Aβ plaques in the *App*^*NL-G-F*^ retina could be due to lower expression levels of Aβ compared to the brain, as shown for other AD mouse models [1], or to different clearance rates. However, importantly, it confers a unique strength to the *App*^*NL-G-F*^ retina as a model for AD research, as it creates a time window to investigate the earliest phases of AD pathogenesis and is closer to the slow disease progression seen in humans. A second asset of the *App*^*NL-G-F*^ retina is that, by diverging from many other available AD models, its phenotype shows striking similarities to the neuropathological changes seen in retinas of human AD patients. This includes the shape and size of the retinal plaques, which we found to be 5-10 times smaller than their counterparts in the brain [22, 23]; the localization of these plaques to the inner retinal layers – where endogenous APP expression has been observed [17, 25] – and associated with retinal blood vessels; as well as the microglial reactivity, astrocytosis and changes in retinal vessel morphology. Of note, the manifestations of retinal vascular amyloidosis are of particular interest in light of the recent findings by Shi *et al*., who reported pericyte loss accompanying retinal vascular amyloidosis in the human AD retina [44]. With one exception [14], this phenotype is thus far unstudied in AD mouse models and our data suggest that the *App*^*NL-G-F*^ retina can be used to further explore this retinal manifestation of AD. Furthermore, the *App*^*NL-G-F*^ retina also appears to be a research model for in-depth investigation of the synaptotoxic effects of Aβ, given the converging data on (soluble) Aβ accumulation in synapses in the inner plexiform layer and electrophysiological abnormalities of amacrine and bipolar cells. Notably, an explanation for the diverging and more severe phenotype of many other transgenic AD mouse models may be sought in the fact that those models are far removed from the human disease state, with non-endogenous promotors, overexpression of APP byproducts, PS1 mutation and/or abnormal APP and Aβ expression levels. This study is pioneering in that it is the first-time retinal phenotyping study of the *App*^*NL- G-F*^ knock-in mouse, which was created to overcome all of the aforementioned artefacts and currently still is unique in its kind. Finally, we have also showcased here the unique potential of the retina for *in vivo* monitoring of structural and functional changes in AD. Of the battery of tests used, hyperspectral imaging and electroretinograms proved to be sensitive read-outs to assess AD disease status. Results from our *post mortem* studies suggest that these may in future studies be complemented with confocal scanning ophthalmoscopy for microgliosis and optical coherence tomography angiography or fundus imaging for detailed retinal vascular assessments. Together, this creates a set of *in vivo* read-outs for preclinical investigation that cannot be matched by any existing *in vivo* brain research methods. Notably, the fact that identical techniques and outcome measures can be used in mice and patients, confers potential translational value to this research.

A better understanding of the time course of the pathological processes underlying preclinical AD, is an important prerequisite for the development of new therapies targeting this phase. Furthermore, retinal imaging may provide an accurate and convenient indication of the efficacy of therapies. Currently, Aβ status is evaluated via CSF assays or PET imaging, which is mainly indicative for the deposition of fibrillary forms of Aβ. As such, these do not always provide information about oligomeric Aβ, which may be the more relevant Aβ species to target [46, 49]. Our data substantiate that hyperspectral imaging is a highly sensitive technique that can be used to quantify Aβ burden in the earliest stages of disease. This imaging spectroscopy technique is based on the fact that soluble Aβ aggregates – most likely Aβ_42_ – interfere with traversing light, leading to Rayleigh light scattering and a characteristic hyperspectral signature, the magnitude of which appear to be proportional to the amount of Aβ present [29, 30]. In the past year, following promising *post mortem* and *in vivo* studies in both animal and human retinas [28–30], results of the first clinical trial showing that *in vivo* retinal hyperspectral imaging can discriminate between Aβ PET-positive cases and controls emerged [13]. This, together with our data, suggests that hyperspectral imaging can be used to monitor retinal Aβ accumulation, with the potential to provide quantitative measures of Aβ oligomers. As such, hyperspectral imaging in the *App*^*NL-G-F*^ retina may become a gateway to improved preclinical studies of the new generation of Aβ therapeutics. Finally, on the long run, this research may also lay the foundations for the rational use of this retinal imaging biomarker for AD diagnosis, disease monitoring, patient stratification and even population-wide screenings.

## V. Conclusion

To conclude, decades of AD research have taught us that we should not try to unravel AD pathogenesis from a simple, neuron-centric perspective, or by assuming a linear series of events leading from Aβ accumulation to dementia. AD is to be understood as a complex pathological cascade, taking decades to gradually evolve from a clinically silent phase to dementia, that is driven by the proteopathic stress imposed by Aβ and Tau conformations, and feedback and feedforward responses of astrocytes, microglia, and vascular cells [48]. Research based on this new framework requires novel approaches, including mouse models that more closely mimic human pathophysiology and allow the study of the earliest, preclinical phases of AD. In this research, we have shown that the retina of *App*^NL-G-F^ mice is a promising model for preclinical AD, in which Aβ oligomers accumulate and pathological cellular responses are elicited. Furthermore, we have added further evidence in support of the potential of hyperspectral imaging to quantify retinal Aβ burden and to serve as an AD biomarker. Altogether, the *App*^NL-G-F^ retina and retinal imaging techniques are unique tools for both fundamental and drug discovery research in the preclinical phase of AD.

## VI.

## List of abbreviations

AD: Alzheimer’s disease
Aβ: amyloid-beta
CNS: central nervous system
ChAT: choline acetyltransferase
GFAP: glial acidic fibrillary protein
GFP: green fluorescent protein
HSI: hyperspectral imaging
Iba1: ionized calcium-binding adapter molecule 1
PFA: paraformaldehyde
RBPMS: RNA-binding protein with multiple splicing
S100B: S100 calcium binding protein B
WT: wild type

## VII. Declarations

### Ethics approval and consent to participate

All animal experiments were performed according to the European directive 2010/63/EU and approved by the KU Leuven institutional ethics committee for animal research.

### Consent for publication

Not applicable

### Availability of data and material

The data sets generated and analyzed during the current study are available from the corresponding author upon reasonable request.

### Authors’ contributions

MV designed the study, performed experiments, analyzed the data and wrote the manuscript; LV, SL, GG performed experiments; XH, LS, JT, LuM analyzed the data; TS and TCS created the *App*^*NL-G-F*^ mice; SL, JT, MJ and LDG implemented the hyperspectral imaging set-up; PDB, IS, BDS, PvW supervised the study and reviewed the manuscript; LM supervised and designed the study, analyzed the data and wrote the Nmanuscript, LDG supervised and designed the study, performed experiments, analyzed the data and wrote the manuscript.

## Acknowledgements

We thank Véronique Brouwers and Evelien Herinckx for their assistance with mouse breeding and experimentation; Wouter Charle, Toon Van Craenendonck and Arnout Standaert for advice on hyperspectral imaging; Isabelle Etienne and Tine Van Bergen (Oxurion, Heverlee, Belgium) for help with immunohistochemistry; RIKEN BioResource Center for creation and distribution of the *App*^*NL-G-*^mice.

## IX. Additional files

**Supplementary figure 1**. Schematic overview of the hyperspectral imaging data analysis. Spectral data from all mice were stacked in a matrix and a corresponding class label (AD or wild type control [CO]) was given for each entry. The between class covariance spectrum was calculated. This spectrum corresponds to the average spectral difference between AD and CO groups. The hyperspectral score for a given spectral pixel was obtained by calculating the dot product between its spectral data and the between class covariance spectrum. The average spectral score for a given mouse was calculated using all pixels of all the HS images collected for that retina.

**Supplementary figure 2**. Electroretinogram measurements in AppNL-G-F mice. Electroretinogram measurements were carried out in both AppNL-G-F and WT mice at 3,6,9,12 and 18 months of age. Data are depicted as mean ± SEM; n=4-12 per group. (pdf file)

