## Supplementary figures for "The *App*^*NL-G-F*^ mouse retina is a site for preclinical Alzheimer’s disease diagnosis and research"

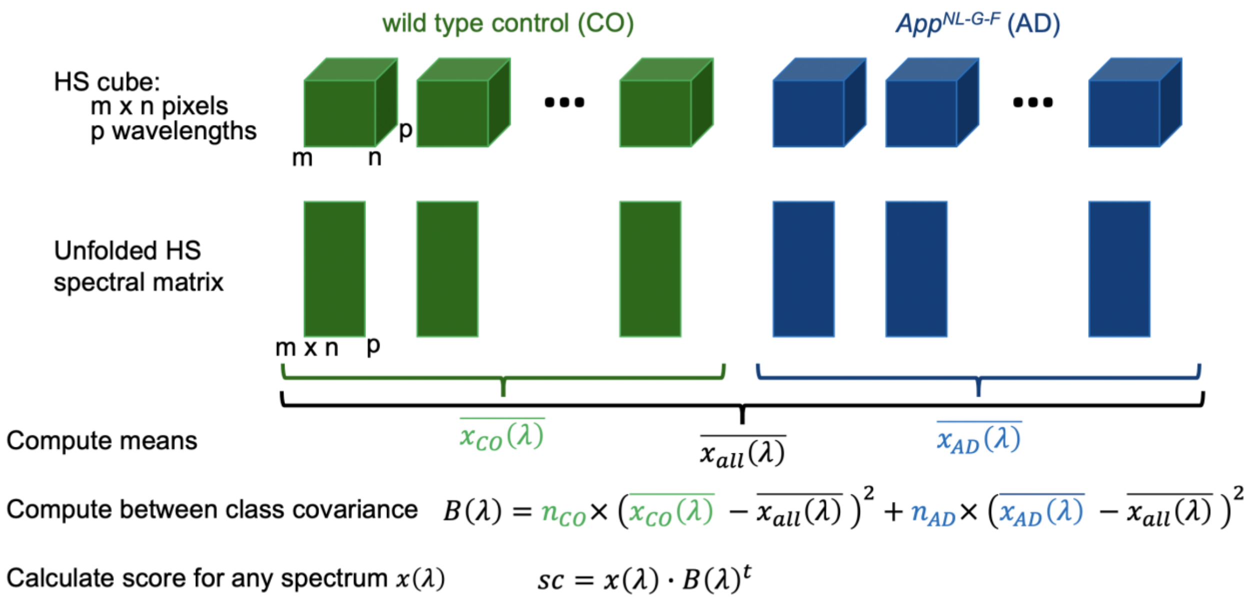


**Supplementary figure 1.** Schematic overview of the hyperspectral imaging data analysis. Spectral data from all mice were stacked in a matrix and a corresponding class label (AD or wild type control [CO]) was given for each entry. The between class covariance spectrum was calculated. This spectrum corresponds to the average spectral difference between AD and CO groups. The hyperspectral score for a given spectral pixel was obtained by calculating the dot product between its spectral data and the between class covariance spectrum. The average spectral score for a given mouse was calculated using all pixels of all the HS images collected for that retina.

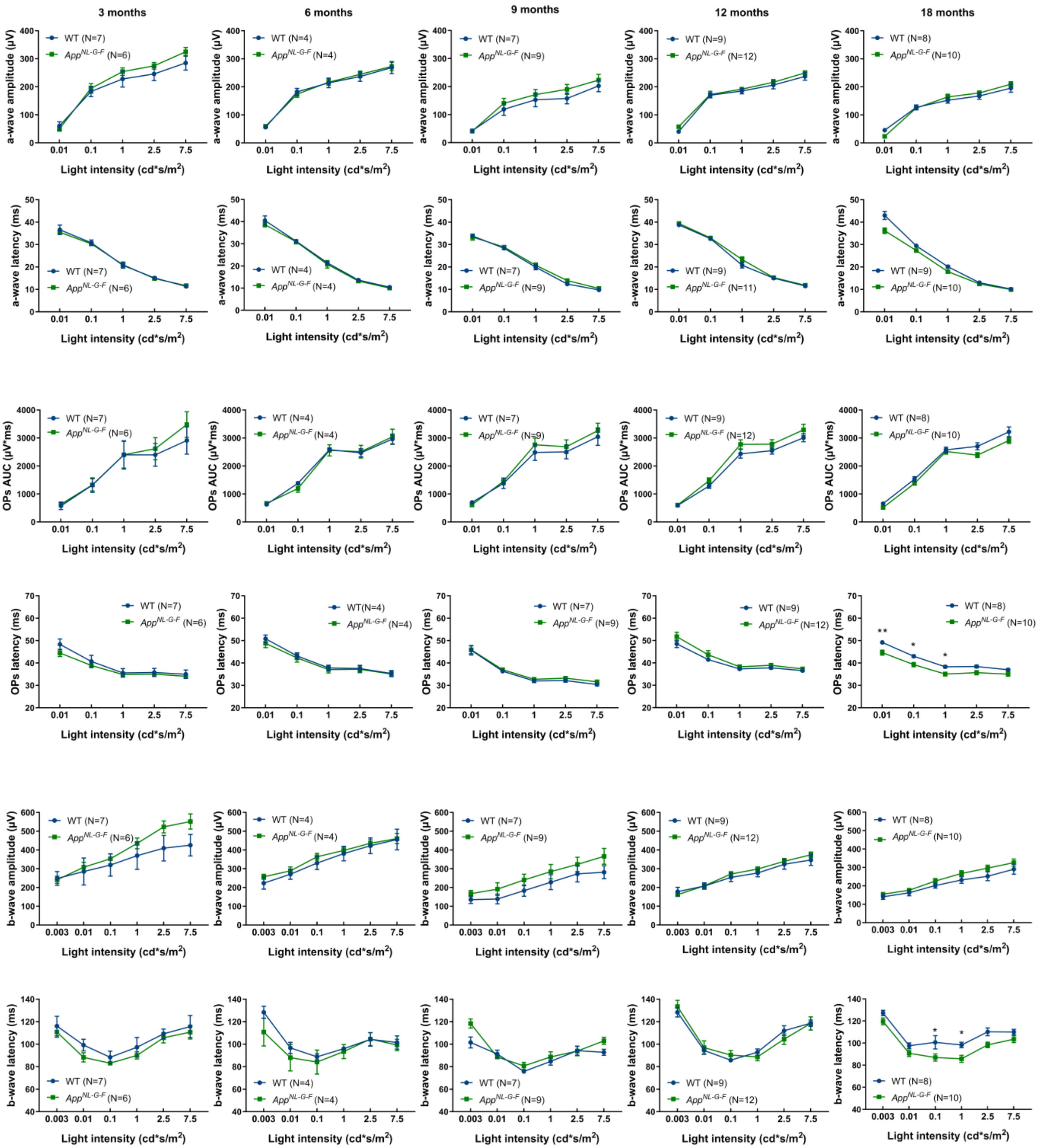
3 months
6 months
9 months
12 months
18 months
a-wave amplitude (μV)
a-wave latency (ms)
OPs AUC (μV*ms)
OPs latency (ms)
b-wave amplitude (μV)
b-wave latency (ms)
Light intensity (cd*s/m²)
WT (N=7)
App^NL-G-F (N=6)
WT (N=4)
App^NL-G-F (N=4)
WT (N=7)
App^NL-G-F (N=9)
WT (N=9)
App^NL-G-F (N=12)
App^NL-G-F (N=11)
WT (N=8)
App^NL-G-F (N=10)
WT (N=9)
WT(N=4)
**
*

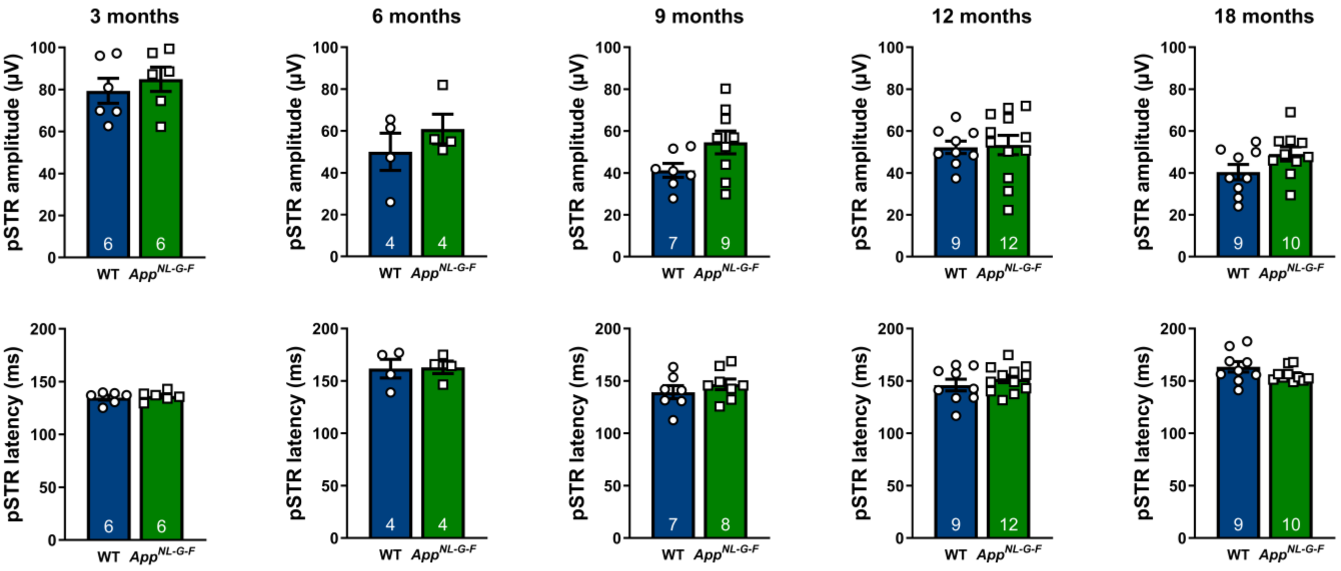


**Supplementary figure 2. Electroretinogram measurements in *App*$^{NL\text{-}G\text{-}F}$ mice.** Electroretinogram measurements were carried out in both *App*$^{NL\text{-}G\text{-}F}$ and WT mice at 3,6,9,12 and 18 months of age. Data are depicted as mean ± SEM; n=4-12 per group.
